# High-throughput multispecies qPCR assays to study the effects of acute thermal stress in three species of *Acipenser* sturgeon

**DOI:** 10.1101/2025.10.11.681731

**Authors:** Hossein Haghighi, Kyle Madden, William S. Bugg, Faith M. Penny, Scott A. Pavey, Nicholas J. Bernier, Kenneth M. Jeffries

## Abstract

In this study, we developed a multispecies OpenArray™ qPCR “chip” to measure the effects of acute thermal stress on the mRNA response of *Acipenser* sturgeons. The qPCR chips were then tested on three species: *Acipenser fulvescens*, *Acipenser oxyrinchus oxyrinchus*, and *Acipenser brevirostrum.* The *A. fulvescens* were from two genetically distinct populations (Burntwood River and Winnipeg River, Manitoba, Canada) that were acclimated to three different temperatures (16, 20, and 24 °C) to quantify the effects of genetic background and acclimation temperatures on the transcriptomic response to acute thermal stress. Additionally, juvenile *A. oxyrinchus oxyrinchus* and *A. brevirostrum* were acclimated to 10 °C and then experienced acute thermal stress. All three species were sampled for gill and liver before and after experiencing acute thermal stress, which was applied using a CT_max_ test. The expression of 52 genes of interest was quantified using the multispecies qPCR chip for all three species, along with 4 reference genes. We showed that acute thermal stress affected the mRNA expression of 17-26 of the thermal metabolic stress genes that were assayed using the chip, depending on the species. However, the mRNA patterns varied between species and genetically distinct populations.

## Introduction

Average water temperatures are increasing due to climate change (Houghton, 2001), which can have profound effects on aquatic environments globally (Fernández, 2022; Ryan & Ryan, 2006). Ectotherms, such as fishes, are sensitive to changes in water temperature (Burraco et al., 2020) because their internal temperature is highly dependent on ambient temperature and temperature directly influences their metabolic rate (Lear et al., 2020). Fishes have evolved to tolerate fluctuations in temperature in their natural distribution (Cuculescu et al., 1998), however, with rapid episodic changes in temperature, such as heat waves, full thermal acclimation is not always possible (Volkoff & Rønnestad, 2020). Thus, rapid temperature increases caused by heat waves have the potential to result in population-level impacts on fishes (Morgan et al., 2020).

The metabolic rate of animals represents the energy needed for maintenance, growth, and reproduction (Riemer et al., 2018). Because temperature determines ectotherms’ metabolic rate, high temperature can increase the cellular energy demand (Sun et al., 2019). In the presence of sufficient oxygen supply to tissues, glucose is metabolized to pyruvate through glycolysis, which is then shuttled into mitochondria where it goes through the tricarboxylic acid (TCA) cycle and is used to produce ATP through oxidative phosphorylation (Bertram et al., 2006). However, in the absence of oxygen, anaerobic glycolysis is used to produce ATP, which has a lower efficiency for producing ATP (Vazquez et al., 2010). In this process, after glycolysis, pyruvate is converted to lactate by lactate dehydrogenase and the by-product of this reaction, nicotinamide adenine dinucleotide (NAD^+^), is used to produce ATP (Chandel, 2021). Changes in the activity of enzymes involved in the aerobic and anaerobic metabolism of glucose have been reported in *Horabagrus brachysoma*, *Sparus aurata*, and *Notothenia rossii* in response to thermal stress (Dalvi et al., 2017; Feidantsis et al., 2009; Forgati et al., 2017). In addition, stress hormones such as catecholamines and cortisol, are important regulators of energy flux in adipose tissue (Peckett et al., 2011). Changes in the expression of genes involved in the metabolism of lipids have also been reported in several species in response to thermal stress (Ma et al., 2015; Balbuena-Pecino et al., 2019; Zhao et al., 2021). Moreover, increased cortisol levels are often associated with situations that result in anaerobic metabolism (Boeck et al., 2001). If not enough ATP is produced to match the consumption of ATP, important activities such as protein production and ion regulation may fail, eventually resulting in cell death (Staples & Buck, 2009).

Acute thermal stress leads to physiological responses in the fish (Morgan et al., 2019), which include changes at the cellular level (Somero, 2020). Upregulation of heat shock proteins (HSP), important in the folding of proteins (Craig et al., 1993; Whitley et al., 1999), is a common cellular response observed in aquatic animals exposed to increased temperature (Zheng et al., 2019). Thermal stress has also been observed to increase the production of reactive oxygen species (ROS) (Slimen et al., 2014). Typically, antioxidant proteins such as superoxide dismutase (SOD), glutathione peroxidase (GPX), and catalase (CAT) detoxify ROS and minimize the adverse effects of these compounds (Matès, 2000). When the production of ROS is more than the antioxidant capacity of cells, ROS can cause damage to cells by lipid peroxidation, and alteration of proteins and DNA, which can lead to cell death (Fleury et al., 2002).

Programmed cell death, or apoptosis, is a type of cell death that can be caused by intracellular stress such as increased levels of ROS, especially through the activation of intrinsic apoptotic pathways (Villalpando-Rodriguez & Gibson, 2021). In this phenomenon, the release of cytochrome c from mitochondria results in the activation of apoptotic protease activating factor-1 (Apaf-1), which aids in the formation of an apoptosome complex that eventually causes the activation of executioner caspases by initiator caspases (Green & Llambi, 2015). Executioner caspases, such as caspase-3, activate cytoplasmic endonucleases and proteases that degrade nucleic acids in the nucleus and proteins (Elmore, 2007), respectively, leading to controlled cell death.

Sturgeons are an ancient group of fishes mostly dwelling in the river systems of the northern hemisphere (Carmona et al., 2009). Since the 1800’s, populations of these species have been threatened by factors such as overfishing and habitat destruction (Lenhardt et al., 2006). As a result, several species of sturgeons are listed as endangered or critically endangered in the International Union for Conservation of Nature (IUCN) red list (IUCN, 2025). Increasing water temperature caused by climate change is one of the threatening factors that affects freshwater fishes’ life from different aspects (Whitney et al., 2016). Consequently, there is a need to understand the effects of temperature on the physiological performance and thermotolerance of fishes (Lefevre et al., 2021). Transcriptomic-based approaches can help us to investigate the physiological status of wild animals, which can connect the transcriptomic profile of animals to fitness related characteristics (Jeffries et al., 2021). Quantitative PCR (qPCR) has been used in several studies on *Acipenser* sturgeons to investigate the cellular and systemic responses to increased temperature (Bugg et al., 2020; Chen et al., 2022; Bugg et al., 2023; Penny et al., 2023; Wang et al., 2023).

The objective of this study was to develop a multispecies *Acipenser* OpenArray™ qPCR “chip” to estimate the expression of 56 genes coding proteins with important functions in the immune system, metabolism, osmoregulation, response to hypoxia, apoptosis, general stress response, and oxidative stress response. We then used this *Acipenser* chip to quantify the effects of acute thermal stress after exposure to a critical thermal maximum trial (CT_max_) protocol during early life stages in three species of *Acipenser* sturgeon: lake sturgeon (*Acipenser fulvescens*), Shortnose sturgeon (*Acipenser brevirostrum*), and Atlantic sturgeon (*Acipenser oxyrinchus oxyrinchus*). These species are endemic to North America and their populations have decreased in recent decades because of anthropogenic activities (Pollock et al., 2015; Fernandes et al., 2010; Kynard & Horgan, 2002). Despite being morphologically similar, and for shortnose and Atlantic sturgeon, existing in similar geographical areas, these three focal species diverged millions of years ago and have different life histories and ploidies (Trifonov et al., 2016; Penny et al., 2023). In this study, the expression of these genes after exposure to acute thermal stress was measured in the gill and liver to test the ability of the *Acipenser* chip to quantify the transcriptomic response to temperature across different species.

## Material and Methods

### Primer and probe optimization for the OpenArray™ chip

We designed a multispecies *Acipenser* OpenArray™ qPCR chip that contains primers and probes for amplifying and detecting 56 genes in *Acipenser* sturgeons. The chip comprises genes involved in the immune system (6 genes), metabolism (15 genes), osmoregulation (4 genes), response to hypoxia (7 genes), apoptosis (4 genes), oxidative stress response (7 genes), general stress response (9 genes), and endogenous controls (4 genes). Each chip can quantify the expression of the 56 genes in 24 samples in duplicate (56 genes × 24 samples × 2 technical replicates = 2688 reactions). The produced cDNA from each sample is loaded on each chip using QuantStudio™ 12K Flex Accufill System (Thermo Fisher Scientific, USA). The chips are run on a QuantStudio 12K Flex Real-Time PCR System (Thermo Fisher Scientific, USA), which can run four chips at a time increasing the number of samples measured each time to 96 (10752 reactions).

To design the universal OpenArray™ chip for *Acipenser* sturgeons, the available genomic and transcriptomic information of different *Acipenser* species was acquired from several sources such as RNA sequencing data of lake sturgeon (Bugg et al., 2023), white sturgeon (*Acipenser transmontanus*) (Doering et al., 2016), Siberian sturgeon (*Acipenser baerii*) (Klopp et al., 2020), and online DNA and RNA sequences belonging to different *Acipenser* species available on National Center for Biotechnology Information (NCBI) (www. https://www.ncbi.nlm.nih.gov/). After the alignment of sequences from different species using Geneious Prime (2023.0.1), primers and probes for each gene were designed based on the consensus sequence at the conserved region of each gene using Primer Express (3.0.1), following a similar protocol used for designing multispecies chips for salmonids (Islam et al., 2024).

The designed primers were tested on lake sturgeon, Atlantic sturgeon, shortnose sturgeon, and white sturgeon with the SYBR Green® detection on a QuantStudio™ 5 Real-Time PCR System (Applied Biosystems, USA). The quality of the amplification and the specificity of primers were checked by monitoring amplification curves and melt curves, respectively. After generating standard curves, the efficiency of PCR reactions was calculated for each primer by the formula E=[10^(1/-slope)^]-1(Wong & Medrano, 2005). Primers with efficiencies between 80 and 120% were used for developing the OpenArray™ chip following established protocols for optimal assay design for the OpenArray technology. The chips were then tested on three species of *Acipenser* sturgeon that underwent CT_max_ protocols outlined below. The sequence of primers and probes and the calculated efficiency for primers are listed in the supplementary Table 1.

## Experimental design

### Lake sturgeon

The tissue samples of *A. fulvescens* used in this study come from a previous study investigating the effects of acclimation temperature on the thermal physiology of two distinct populations from Burntwood River and Winnipeg River in northern and southern Manitoba, respectively (Bugg et al., 2020). The juvenile lake sturgeons were produced by fertilizing gametes harvested from wild-caught females and males from the two populations at the University of Manitoba, Canada. Beginning at 30 days post fertilization (DPF), the juveniles from these two populations were acclimated to 16, 20, and 24 °C for 30 days. After 30 days (60 DPF), and before the (CT_max_) trial (Pre-CT_max_), eight fish from each acclimation temperature of both Winnipeg River (1.02 ± 0.41 g; 67.71 ± 10 mm *L*_T_) and Burntwood River (1.14 ± 0.51 g; 71.51 ± 10 mm *L*_T_) were euthanized by immersion in an overdose of tricaine methane sulfonate solution buffered with an equal volume of sodium bicarbonate. The gill tissue was collected and preserved in RNA*later®* (Thermo Fisher Scientific, Waltham, USA), until transferred to −80 °C for storage. Another eight fish from each acclimation temperature were netted and placed individually in well-aerated experimental units (∼200 ml of water volume and 9.5 cm long × 5 cm across) at the acclimation temperature one hour before the CT_max_ trial. The CT_max_ experiment was conducted by increasing the water temperature in the tank by 0.3 °C min^−1^ with constant aeration until the fish could not regain proper orientation after a physical disturbance. At this moment, the fish was euthanized by immersing in tricaine methanesulfonate solution (250 mg.L^-1^) buffered with an equal volume of sodium bicarbonate, and the gill tissue was removed and preserved.

### Atlantic and shortnose sturgeon

Samples of Atlantic and shortnose sturgeon came from another study investigating the physiological response to acute thermal stress (Penny et al., 2023). In that study, two-year-old juvenile Atlantic sturgeon (122 ± 10. 3 g; 33.8 ± 0.84 cm total length (*L*_T_)) and shortnose sturgeon (136 ± 8.6 g; 32.9 ± 1.0 cm *L*_T_), were obtained from Acadian sturgeon and Caviar (Carter’s Point, Kingston, New Brunswick, Canada) and brought to the University of New Brunswick, Canada. Fish from both species were separated into two groups: the CT_max_ group and the Control group. They were then acclimated to 10 °C for one month. The CT_max_ was performed on the CT_max_ groups by increasing temperature at a rate of 0.1 °C min^−1^ until the fish was not able to maintain an upright orientation. At this timepoint, the fish was removed from the tank, and after anesthetizing in a tricaine methanesulphonate solution (MS-222; 1 g⋅l^-1^) buffered in Sodium bicarbonate (2 g⋅l^-1^) and euthanizing by cervical dislocation, the gill and liver tissues were collected. Eight fish from the Control group, which experienced the same handling process except for no increase in temperature, were collected and sampled the same way. Samples from both groups were preserved in RNA*later*® and stored at −80 °C.

### RNA extraction, cDNA synthesis

Total RNA from liver and gill samples of Atlantic and shortnose sturgeon, and liver tissue of white sturgeon, was extracted using RNeasy® Plus Micro Kit (QIAGEN, Germany) and PureLink RNA Mini Kits (Invitrogen; Ambion Life Technologies) was used for the lake sturgeon gill samples. Samples from all three species were homogenized using a TissueLyser (QIAGEN) at 50 Hz for 5 min in the lysis buffer available in each specific kit, and RNA purification was done following the manufacturer’s instructions. NanoDrop One (Thermo Fisher Scientific) was used to measure RNA concentration and purity. To measure the integrity of RNA, 400 ng of RNA from each sample was loaded on 1% agarose gel to visually check 18S rRNA and 28S rRNA bands. Before cDNA synthesis, genomic DNA was eliminated by treating 1000 ng of total RNA with Invitrogen™ *ezDNase* enzyme (Thermo Fisher Scientific) at 37 °C for 2 minutes. The DNase-treated RNA was used directly for cDNA synthesis using QuantiTect® Reverse Transcription Kit (QIAGEN) following the manufacturer’s instructions.

### Relative gene expression analysis

To quantify the expression of genes under study using the universal *Acipenser* OpenArray™ chip, the synthesized cDNA was diluted 1:4 with nuclease-free water. Then, 2.5 µl of the diluted cDNA was mixed with 2.5 µl of 2X TaqMan Real-Time PCR Master Mix (Thermo Fisher Scientific, USA). The samples were loaded onto the chip in duplicate using QuantStudio™ 12K Flex Accufill System and real-time PCR was done using QuantStudio 12K Flex Real-Time PCR System following the manufacturer’s instruction. The quality of amplification was assessed using the QuantStudio 12K Flex Real-Time PCR System software and samples with sigmoidal amplification curves and amplification scores higher than 0.125 were selected for further analysis. To calculate the efficiency of PCR for each gene, baseline-corrected fluorescence data was exported from the software, and the efficiency of each reaction was calculated by LinRegPCR (Version 2021.2). For each gene, the mean value of all the efficiencies from single reactions was calculated for each species and listed in Supplementary Table 2. The average efficiency value was used to calculate the relative gene expression by applying the Pfaffl Method (Pfaffl, 2001). To measure relative gene expression in lake sturgeon, all the treatments for each population were normalized to the samples collected from treatment acclimated to 16 °C before the CT_max_ trial (i.e., Pre-16). For Atlantic and shortnose sturgeon, the expression of each gene in the CT_max_ group was calculated relative to the individual species’ Control group. Some gene expression data were excluded from the relative expression analysis due to low PCR efficiency. Specifically, data for *cam1*, *mhc1a*, and *atp1a* were removed for all species. Also, *casp3* was excluded for lake sturgeons, the liver of Atlantic sturgeon, and the gills of shortnose sturgeon. In Atlantic sturgeon, *tp53*, *cox1*, *lipe*, and *ca15* were removed from liver data, while *lipe*, *tp53*, and *cox1* were also excluded from gill data. Additionally, *ca15* was excluded from the liver data of shortnose sturgeon.

### Statistical analysis

All the statistical analyses were conducted using R 4.3.2 (R Core Team, 2024). Shapiro– Wilks’s tests were used to assess the normality of data distribution in treatment groups for all three species. Because the relative gene expression for individuals was not normally distributed, non-parametric statistical methods were used to test differences between groups (p-value < 0.05) in gene expression. Kruskal-Wallis multiple comparison tests followed by a Dunn’s post-hoc test was performed using ‘MultNonParam’ and ‘dunn.test’ packages to compare results across treatments in lake sturgeon. Then, a Bonferroni corrected p-value of 0.0033 was used to determine statistical significance. For Atlantic and shortnose sturgeon, the comparison between groups was conducted using Mann–Whitney U tests. To check the overall transcriptomic change in response to thermal stress, unsupervised Principal Component Analysis (PCA) was conducted for each species using C_t_ values normalized to the four internal control genes.

## Results

### Lake sturgeon

The CT_max_ measured for the two populations of lake sturgeon was slightly higher in the Winnipeg River fish than in the northern Burntwood River fish (Table 1). After CT_max_, the *hsp* genes exhibited the highest upregulation in both lake sturgeon populations (Fig. 1A; 2A). The *hsp70a* gene showed marked upregulation following CT_max_ across all acclimation temperatures in both populations (from 323 to 637 fold change). Similarly, *hsp90a* expression increased significantly in the gill tissue of Burntwood River sturgeons acclimated to 16 (19 fold), 20 (28 fold), and 24 °C (25 fold). In contrast, in the Winnipeg River population, significant upregulation of *hsp90a* was only observed at 20 °C (7.6 fold) and 24 °C (8.8 fold). Increasing acclimation temperature also upregulated *hspa4a* expression after CT_max_ in both populations. However, this effect was statistically significant at 20 °C in the Burntwood River population (2.73 fold), and at 24 °C in the Winnipeg River population (1. 37 fold). Finally, in the Winnipeg River population, the gene coding for cold-inducible RNA binding protein (CIRBP) was significantly downregulated (0.35 fold) after CT_max_ in fish acclimated to 16 °C.

**Fig. 1.**
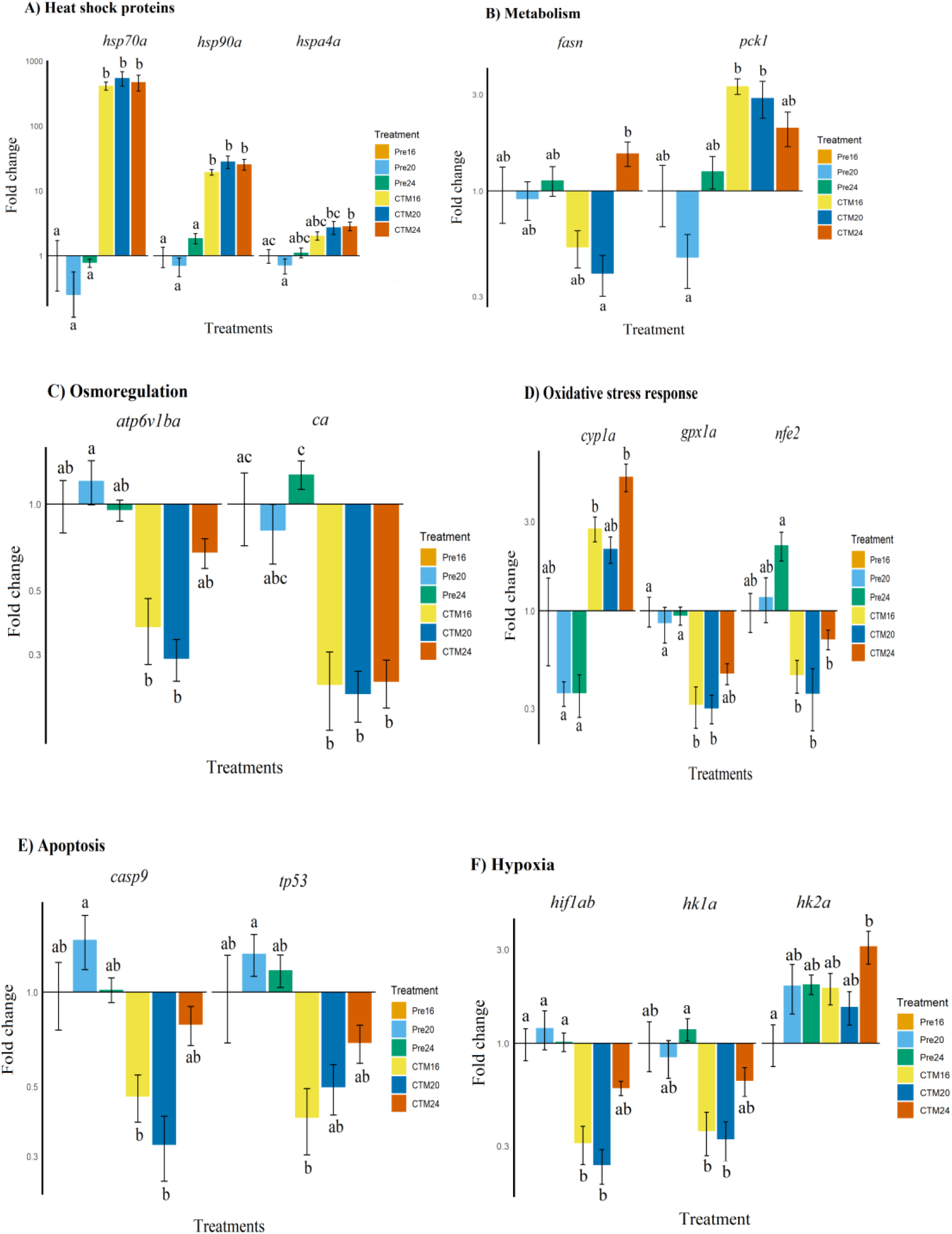
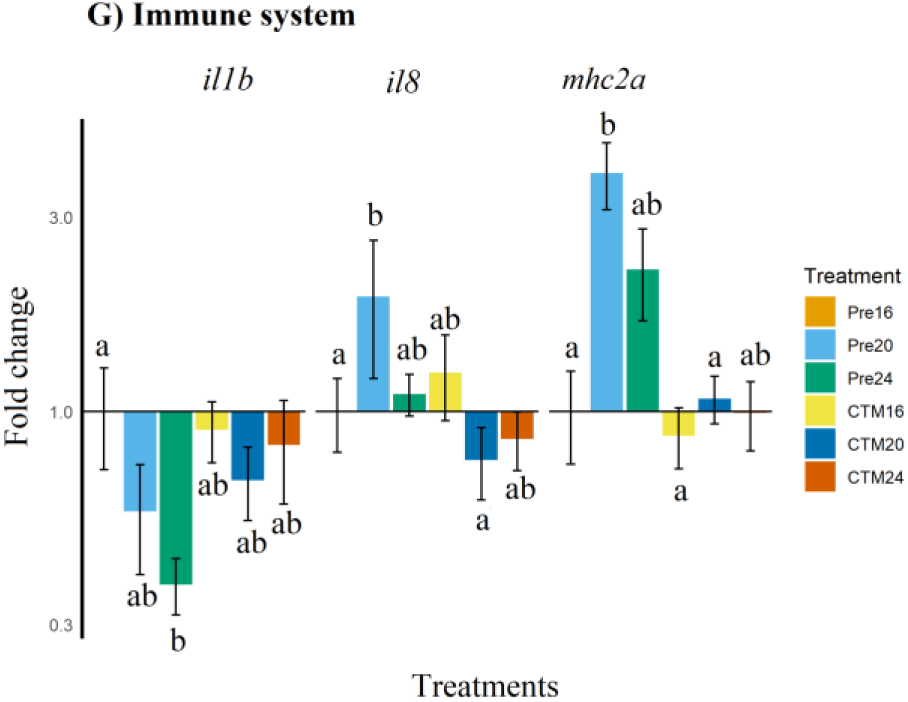
The relative expression of genes with significant differences between groups in the Burntwood River lake sturgeon (*Acipenser fulvescens*) population that were acclimated to 16, 20 and 24 °C. A subset of those fish were then exposed to a CT_max_ trial. Genes involved in different biological processes are presented in different panels. The expression of each gene is relative to the 16 °C acclimation temperature, and groups that do not share a common letter are significantly different from each other. A significant difference between groups was measured by Kruskal–Wallis multiple comparisons with P-values adjusted with the Bonferroni method. For better representation, a logarithmic scale has been used on the y-axis.

**Fig. 2.**
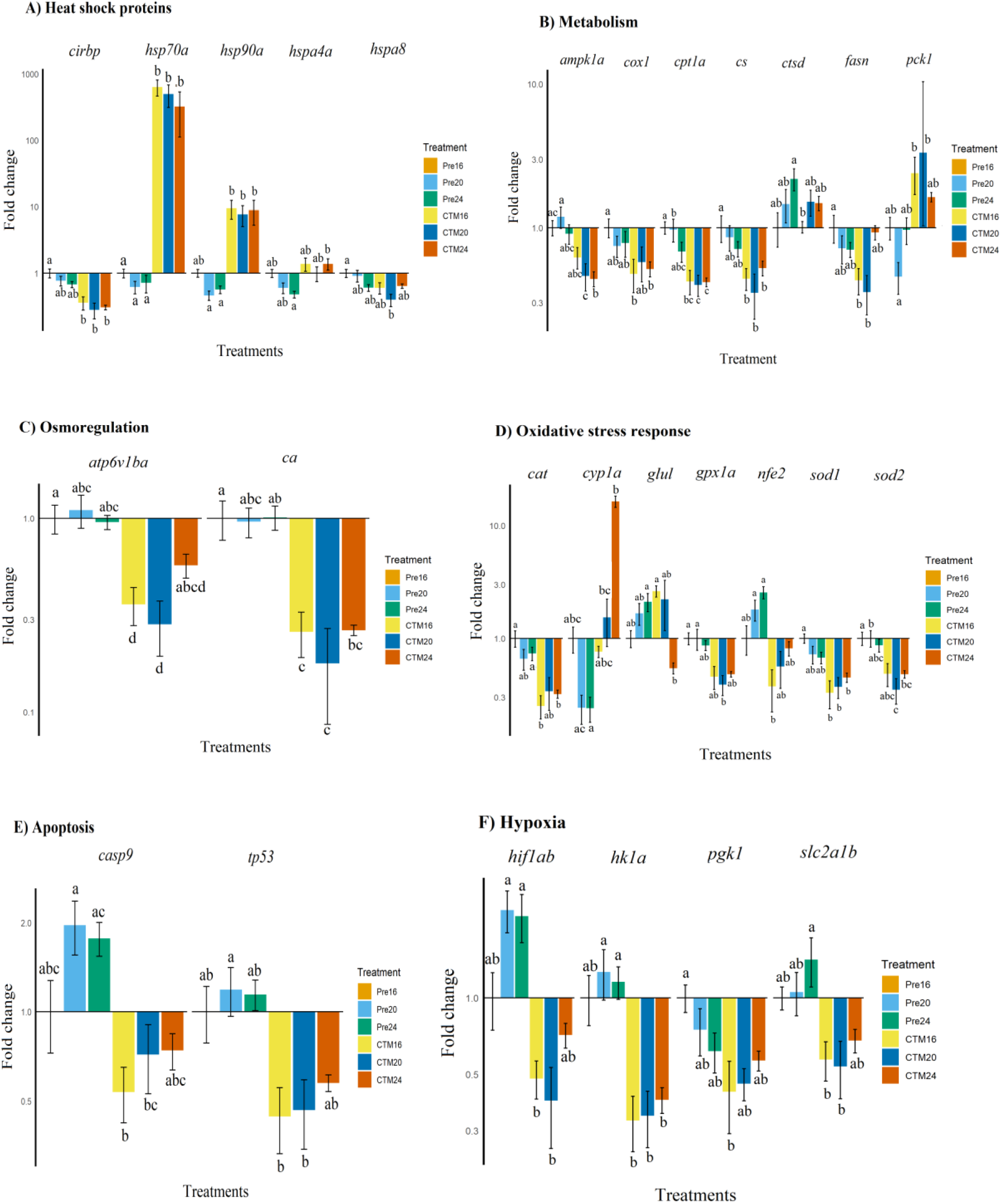
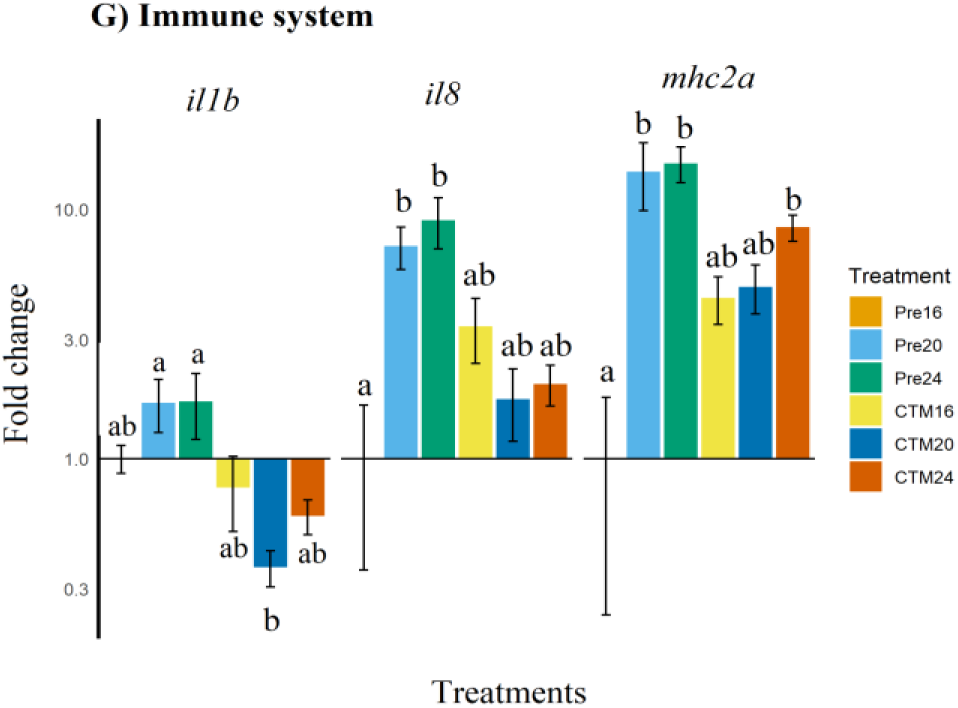
The relative expression of genes with significant differences between groups in the Winnipeg River lake sturgeon (*Acipenser fulvescens*) population that were acclimated to 16, 20, and 24 °C. A subset of those fish were then exposed to a CT_max_ trial. Genes involved in different biological processes are presented in different panels. The expression of each gene is relative to the 16 °C acclimation temperature, and groups that do not share a common letter are significantly different from each other. A significant difference between groups was measured by Kruskal–Wallis multiple comparisons with P-values adjusted with the Bonferroni method. For better representation, a logarithmic scale has been used on the y-axis.

**Table 1.** CT_max_ data of lake sturgeon from Burntwood River and Winnipeg River, Atlantic sturgeon, and shortnose sturgeon. Sixty-day-old lake sturgeon from Burntwood River and Winnipeg River were acclimated to 16, 20, and 24 °C for 30 days and then experienced CT_max_. Two-year-old Atlantic and shortnose sturgeons were acclimated to 10 °C for 30 days, and then experienced CT_max_. Lake sturgeon CT_max_ data are from Bugg et al. 2020. Atlantic sturgeon and shortnose sturgeon CTmax data are from Penny et al. 2023.

| Species, population | Age | Acclimation temperature | Mean CT<sub>max</sub>(°C) + SE |
| --- | --- | --- | --- |
| Lake Sturgeon, Burntwood River | 60 days | 16 °C | 32.21 + 0.13 |
| Lake Sturgeon, Burntwood River | 60 days | 20 °C | 33.90 + 0.15 |
| Lake Sturgeon, Burntwood River | 60 days | 24 °C | 34.74 + 0.24 |
| Lake Sturgeon, Winnipeg River | 60 days | 16 °C | 32.33 + 0.07 |
| Lake Sturgeon, Winnipeg River | 60 days | 20 °C | 34.62 + 0.1 |
| Lake Sturgeon, Winnipeg River | 60 days | 24 °C | 35.48 + 0.24 |
| Atlantic Sturgeon | 2 years | 10 °C | 28.56 + 0.14 |
| Shortnose Sturgeon | 2 years | 10 °C | 27.91 + 0.76 |

Genes involved in energy metabolism were also affected by acute thermal stress, but the response varied by population and acclimation temperature (Fig. 1B; 2B). The *pck1* gene was significantly upregulated in both populations of lake sturgeons acclimated to 20 °C (2.88 and 3.33 fold change in the Burntwood River population and Winnipeg River population, respectively). In contrast, after CT_max_ *cs* (0.44 fold), *fasn* (0.42 fold), and *cox1* (0.47 fold) showed significant downregulation only in the Winnipeg River sturgeons acclimated to 16 °C. Additionally, expressions of *cpt1a* (0.39 fold) and *ampk1a* (0.46 fold) were significantly reduced in response to CT_max_ in the Winnipeg River sturgeons acclimated to 20 °C following CT_max_.

Genes involved in osmoregulation also showed distinct expression patterns in response to acute thermal stress in two populations of lake sturgeons (Fig. 1C; 2C). Expression of *ca* was significantly downregulated (0.3 fold) in Winnipeg River lake sturgeons acclimated to 16 °C, while in the Burntwood River population, this downregulation was only significant at 24 °C (0.3 fold). Similarly, *atp6v1ba* expression was significantly reduced in the Winnipeg River population at 16 °C (0.20 fold) and 20 °C (0.35 fold), but in Burntwood River sturgeon, significant downregulation was observed only at 20 °C (0.28 fold).

Acute thermal stress affected the expression of genes involved in oxidative stress response and xenobiotic metabolism (Fig. 1D; 2D). The expression of *cyp1a* was significantly upregulated in both lake sturgeon populations acclimated to 24 °C (5.2 and 16 fold change in Burntwood River population and Winnipeg River population, respectively). The expression of *nfe2*, a master regulator of antioxidant gene expression, was downregulated after CT_max_ in both populations of lake sturgeons in this study, but this change was only statistically significant in the lake sturgeon from the Burntwood River population acclimated to 24 °C (2 fold). Among the antioxidant genes, *gpx1a* was significantly downregulated in the Burntwood River population acclimated to 16 (0.3 fold) and 20 °C (0.29 fold). In the Winnipeg River population, more antioxidant genes showed significant downregulation; fish acclimated to 16 °C showed reduced expression of *sod1* (0.32 fold) and *cat* (0.25 fold); at 20 °C, *sod2* (0.35 fold) and *gpx1a* (0.38 fold) were downregulated; and at 24 °C, *glul* expression was significantly reduced (0.54 fold). Genes involved in the regulation of apoptosis were also downregulated in response to acute thermal stress; however, only *casp9* showed statistically significant downregulation in fish acclimated to 20 °C in both populations (0.71 fold in the Burntwood River population and 0.32 fold in the Winnipeg River population) (Fig. 3; 4). Genes involved in hypoxia response and glycolysis were mostly downregulated following acute thermal stress (Fig. 3; 4). This downregulation was statistically significant for *hif1ab* in both lake sturgeon populations acclimated to 20 °C (0.39 fold in the Burntwood River population and 0.23 fold downregulation in the Winnipeg River population). Additionally, *hk1a* expression was significantly reduced in the Winnipeg River lake sturgeons acclimated to 20 (0.34 fold) and 24 °C (39 fold).

**Fig. 3.**
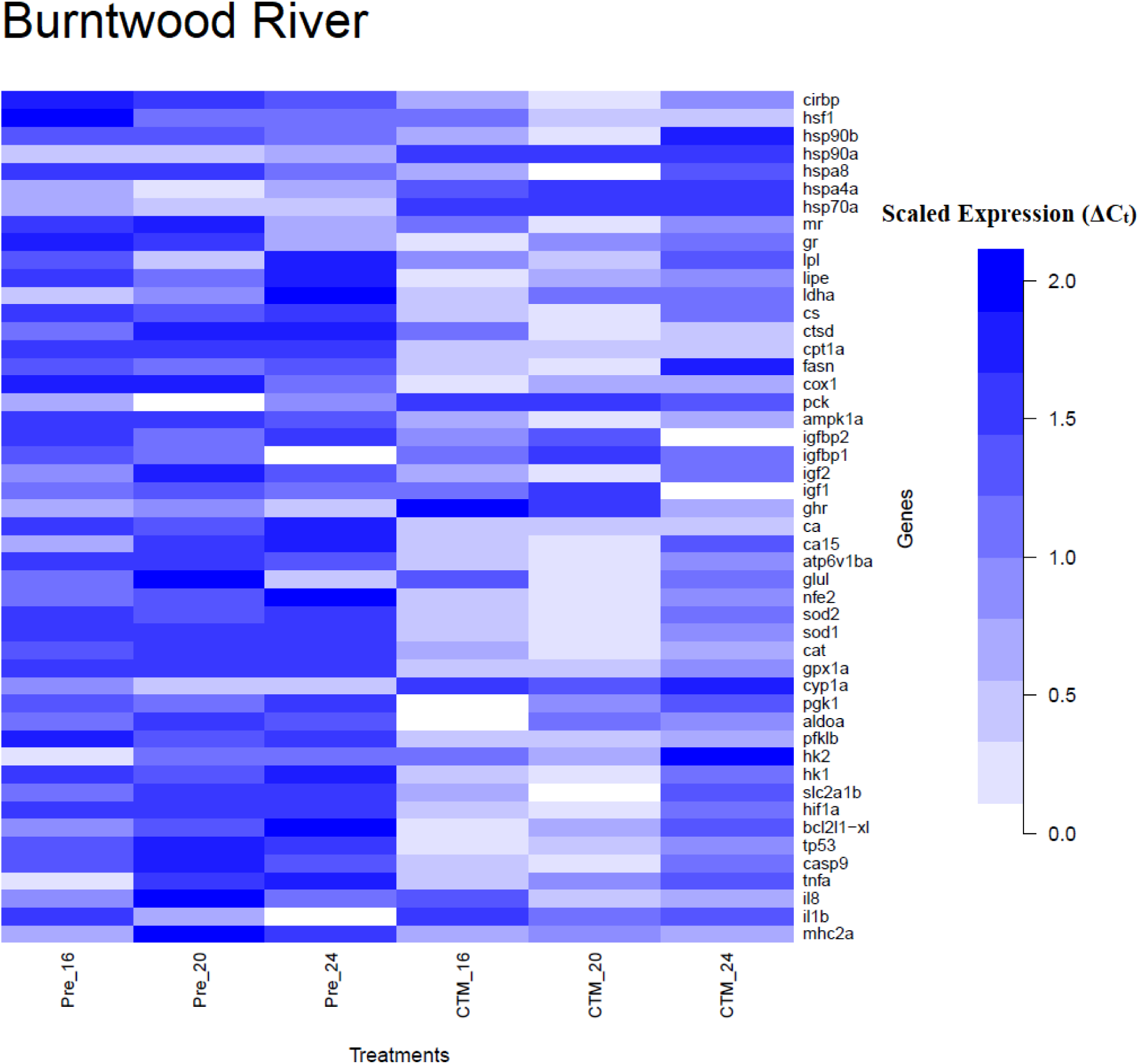
Expression pattern of 52 genes measured in lake sturgeons (*Acipenser fulvescens*) from the Burntwood River population in response to acclimation temperatures and acute thermal stress. Juvenile lake sturgeon were acclimated to 16, 20, and 24 °C for 30 days, and then a subset of them experienced CT_max_. To generate this plot, the expression of each gene in the gill is measured using qPCR, and C_t_ values are normalized to the mean C_t_ value of four control genes and scaled for better representation.

**Fig. 4.**
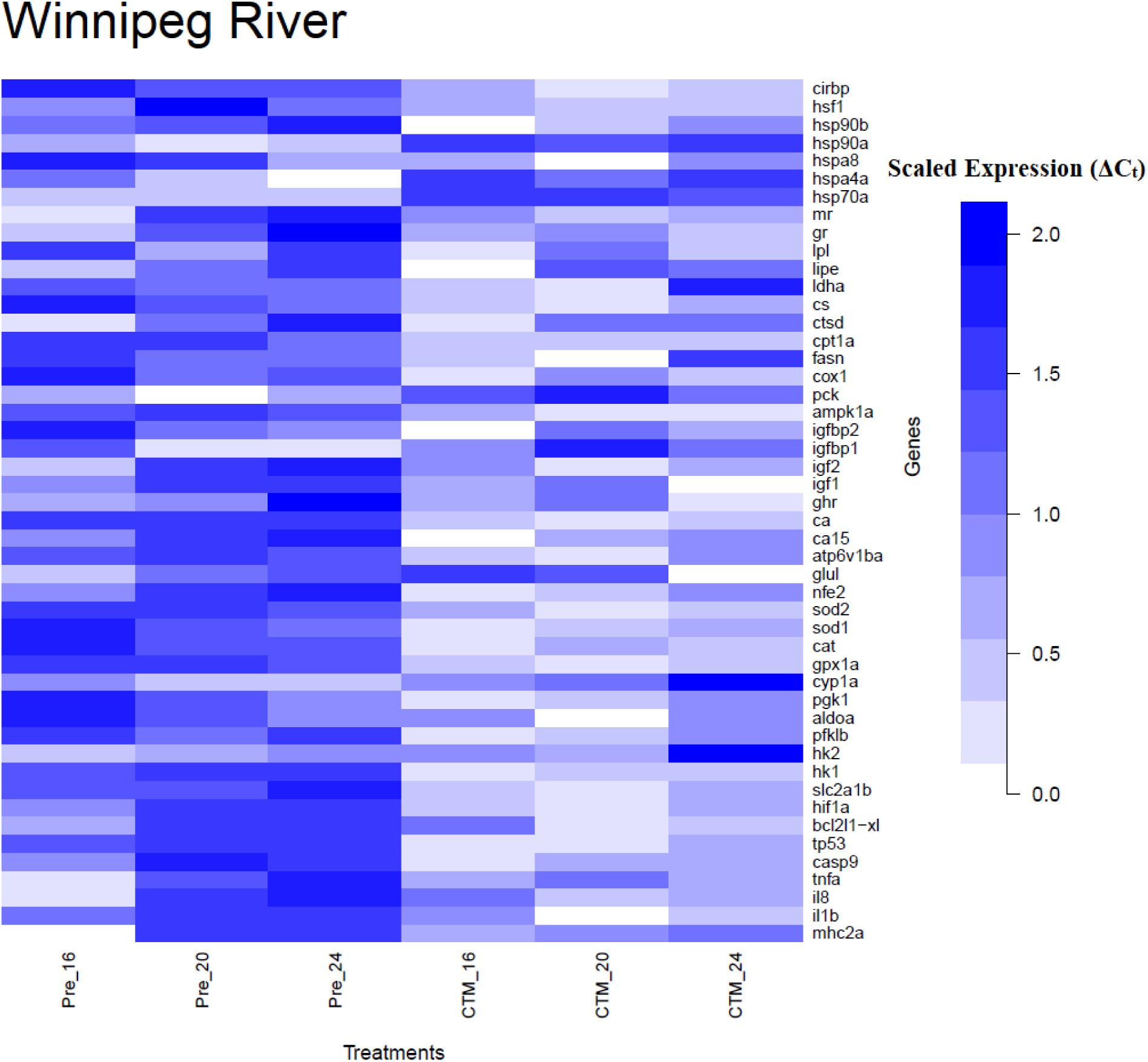
Expression pattern of 52 genes measured in lake sturgeon (*Acipenser fulvescens*) from the Winnipeg River population in response to acclimation temperatures and acute thermal stress. Juvenile lake sturgeon were acclimated to 16, 20, and 24 °C for 30 days, and then a subset of them experienced CT_max_. To generate this plot, the expression of each gene in the gill is measured using qPCR, and C_t_ values are normalized to the mean C_t_ value of four control genes and scaled for better representation.

The effects of acute thermal stress on the immune system were more pronounced in the expression of proinflammatory genes. In the Burntwood River population, *il8* expression was significantly decreased (0.75 fold) after CT_max_ in the fish acclimated to 20 °C, while in the Winnipeg River population, *il1b* was downregulated (0.36 fold) in the same acclimation temperature. Additionally, *mhc2a* expression was significantly downregulated (3.8 fold) in response to CT_max_ in Burntwood River sturgeon acclimated to 20 °C.

### Atlantic and shortnose sturgeon

The CT_max_ values measured in Atlantic and shortnose sturgeon showed slightly higher thermotolerance in Atlantic sturgeon (Table 1). Acute thermal stress significantly upregulated the expression of *hsp70a* in both gill (1476 to 321 fold-changes, respectively) and liver (17 to 73 fold-changes, respectively) tissue of Atlantic and shortnose sturgeon (Fig. 5). Similarly, *hsp90a* was upregulated in the gill tissue of both species (12.5-12 fold) and the liver tissue of shortnose sturgeon (73 fold). In addition, *hspa4a* expression increased in the gill tissue of shortnose sturgeon (2 fold). Following CT_max_, *cirbp* expression was downregulated in the gill tissue of Atlantic sturgeon (0.8 fold), but upregulated in the liver tissue of the shortnose sturgeon (1.3 fold).

**Fig. 5.**
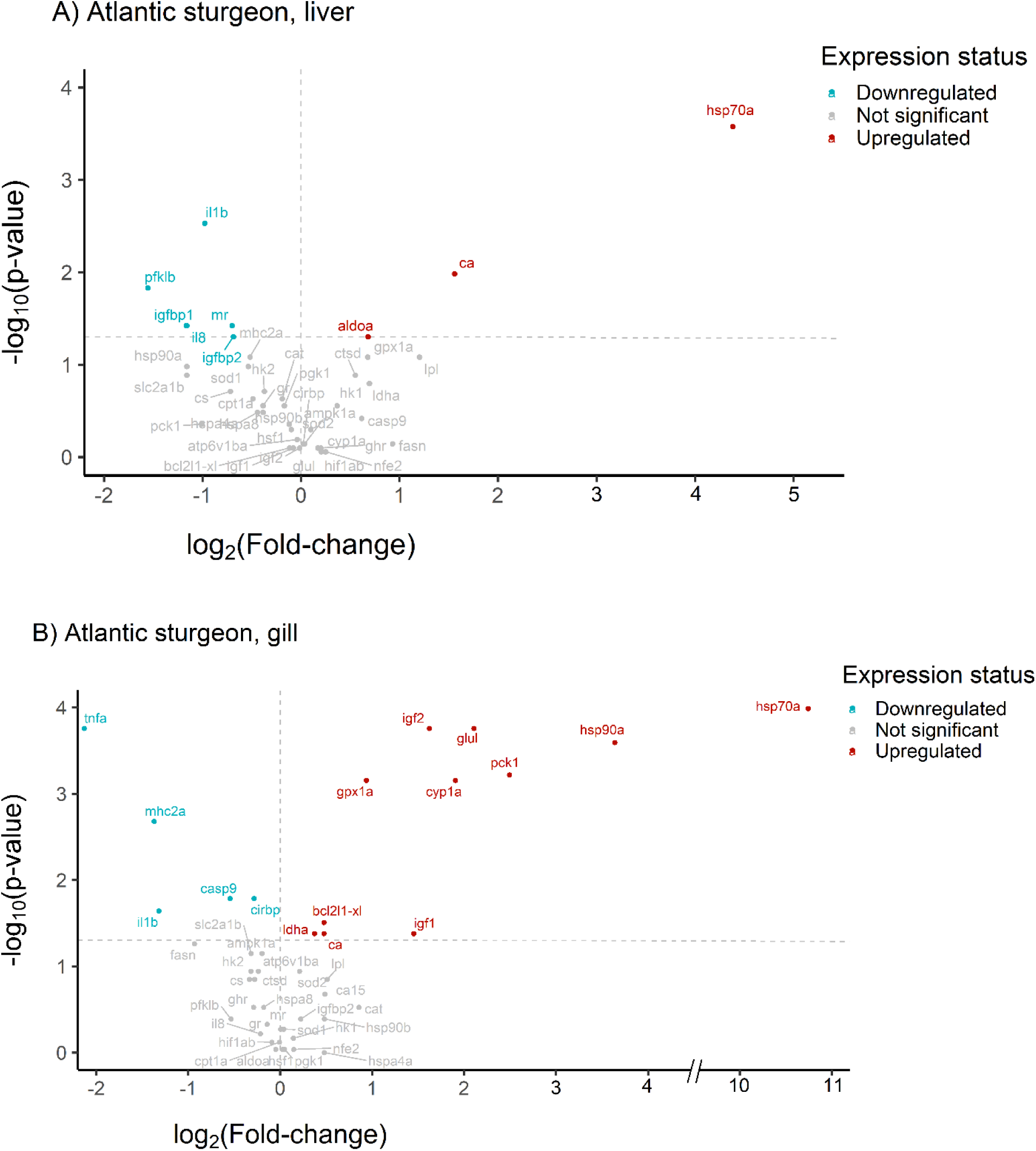
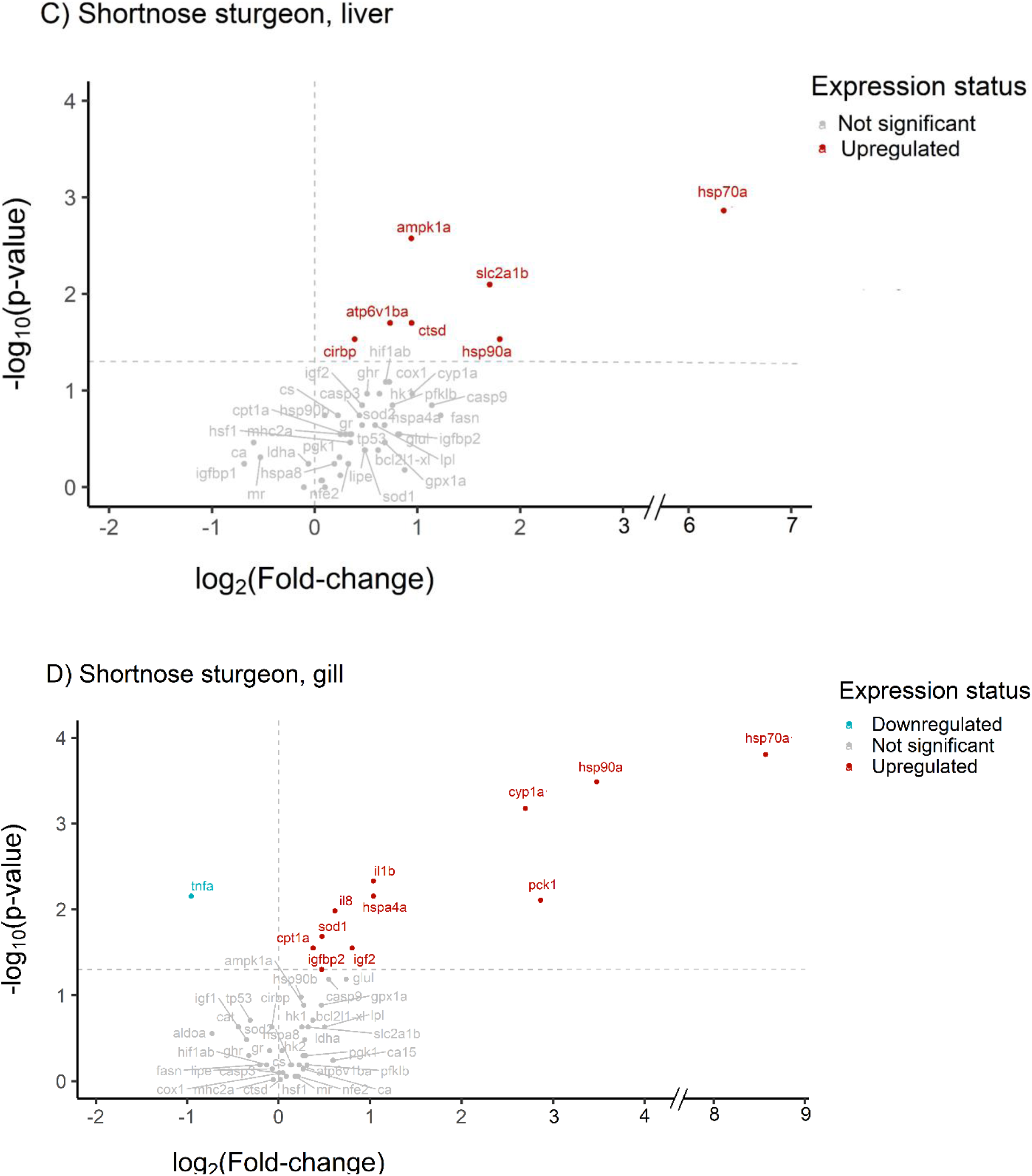
Relative expression of 52 target genes in the liver and gill of Atlantic and shortnose sturgeon, *Acipenser oxyrinchus oxyrinchus, and Acipenser brevirostrum*, in response to CT_max_. Each species was separately acclimated to 10 °C and then experienced CT_max_. The expression of each gene after CT_max_ is calculated relative to a subset of fish before CT_max_, ie., the Control group. The statistical significance between the Control group and CT_max_ group was determined by the Mann–Whitney U test. For better representation, p-values and fold-change expression have been transformed to log10 and log2, respectively. The horizontal dotted line shows a p-value of 0.05, and the vertical dotted line shows a one-time fold-change in expression.

In the Atlantic sturgeon, acute thermal stress significantly downregulated *igfbp1* (0.44 fold) and *igfbp2* (0.62 fold) in the liver, while upregulating *igf1* (2.7 fold), *igf2* (3 fold), *ldha* (1.29 fold), and *pck1* (6.9 fold) in the gill (Fig. 5A; 5B). In shortnose sturgeon, acute thermal stress significantly increased the expression of *cpt1a* (1.29 fold), *pck1* (8 fold), *igf2* (1.74 fold), and *igfbp2* (1.38 fold) in the gill, as well as *ampk1a* (1.91 fold), *ctsd* (1.92 fold) in the liver (Fig. 5C; 5D). In Atlantic sturgeon, *ca* was significantly upregulated in both gill (1.39 fold) and liver (2.95 fold) tissue. In shortnose sturgeon, *atp6v1ba* expression was significantly increased (1.66 fold) in the liver (Fig. 5C). No osmoregulatory genes showed significant changes in expression in the gill tissue of shortnose sturgeon.

The effects of acute thermal stress on the expression of genes associated with xenobiotic metabolism and oxidative stress response were more pronounced in the gill than in the liver (Fig. 5). Similar to observations in lake sturgeon, acute thermal stress increased the expression of *cyp1a* in the gill tissue of Atlantic (3.75 fold) and shortnose sturgeons (7.26 fold) (Fig. 5B; 5D). In Atlantic sturgeon, acute thermal stress significantly upregulated *glul* (4.3 fold) and *gpx1a* (1.9 fold) in the gill, whereas in shortnose sturgeon, only *sod1* (1.38 fold) was upregulated following exposure to CT_max_. In contrast, acute thermal stress did not affect the expression of genes involved in xenobiotic metabolism and oxidative stress response in the liver tissue of either species (Fig. 5A; 5C).

In Atlantic sturgeon, *casp9* expression in the gill was significantly downregulated after CT_max_ (0.68 fold) (Fig. 5B). Conversely, *bcl2l1-xl* was significantly (1.39 fold) upregulated in the gill in response to acute thermal stress. Expression of apoptosis-related genes did not show any significant change in the liver of Atlantic sturgeon or in either the gill or liver tissue of shortnose sturgeon (Fig. 5A; 5C; 5D).

Genes involved in glycolysis and anaerobic metabolism were exclusively upregulated in the liver of Atlantic and shortnose sturgeons. In the Atlantic sturgeon, acute thermal stress significantly upregulated (1.6 fold) the expression of *aldoa*, whereas in shortnose sturgeon, *slc2a1b* was significantly (3.25 fold) upregulated (Fig. 5A; 5C).

In the gill tissue of Atlantic sturgeon, acute thermal stress significantly downregulated the expression of the immune genes *tnfa* (0.22 fold), *mhc2a* (0.38 fold), and *il1b* (0.4 fold), while in the liver, *il1b* (0.5 fold) and *il8* (0.44 fold) were downregulated (Fig. 5A; 5B). In the shortnose sturgeon, the only immune gene significantly affected by CT_max_ was *tnfa,* which was downregulated (0.51 fold) in the gill (Fig. 5C; 5D).

### Principal Component Analysis (PCA)

Principal component analysis (PCA) was used to investigate the overall changes in all the gene expression assayed in response to acute thermal stress and determine the main drivers of differences in gene expression patterns. Principal component analysis revealed that acute thermal stress altered overall gene expression patterns across all three species and both tissues examined (Fig. 6). In both lake sturgeon populations, acclimating to different temperatures changed the expression pattern of 48 genes before and after acute thermal stress (Fig. 6A1; 6B1). Dimension 1 accounted for about 36.1-58 % variation among samples in all PCA plots (Fig. 6). The contribution of genes in the second dimension was aligned more closely with the differential gene expression between temperature treatments in the gill tissues of lake sturgeon, the liver and gill tissue of Atlantic sturgeon, and the gill tissue of shortnose sturgeon (Fig. 6A2; 6B2; 6C2; 6D2; 6F2). In contrast, in the liver tissue of shortnose sturgeon, the contribution of genes in Dimension 1 more closely reflected the differential gene expression pattern between treatments (Fig. 6E).

**Fig. 6.**
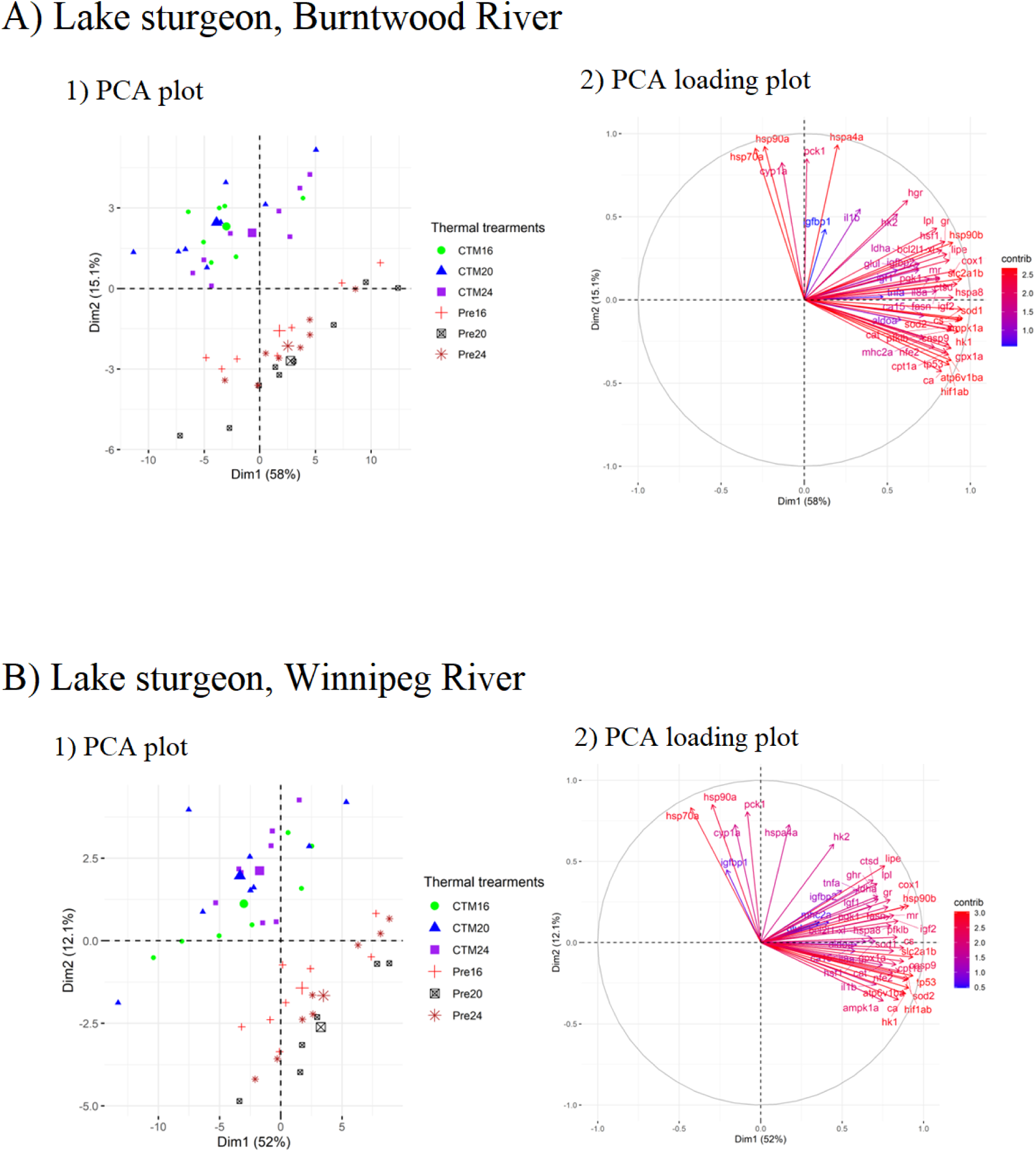
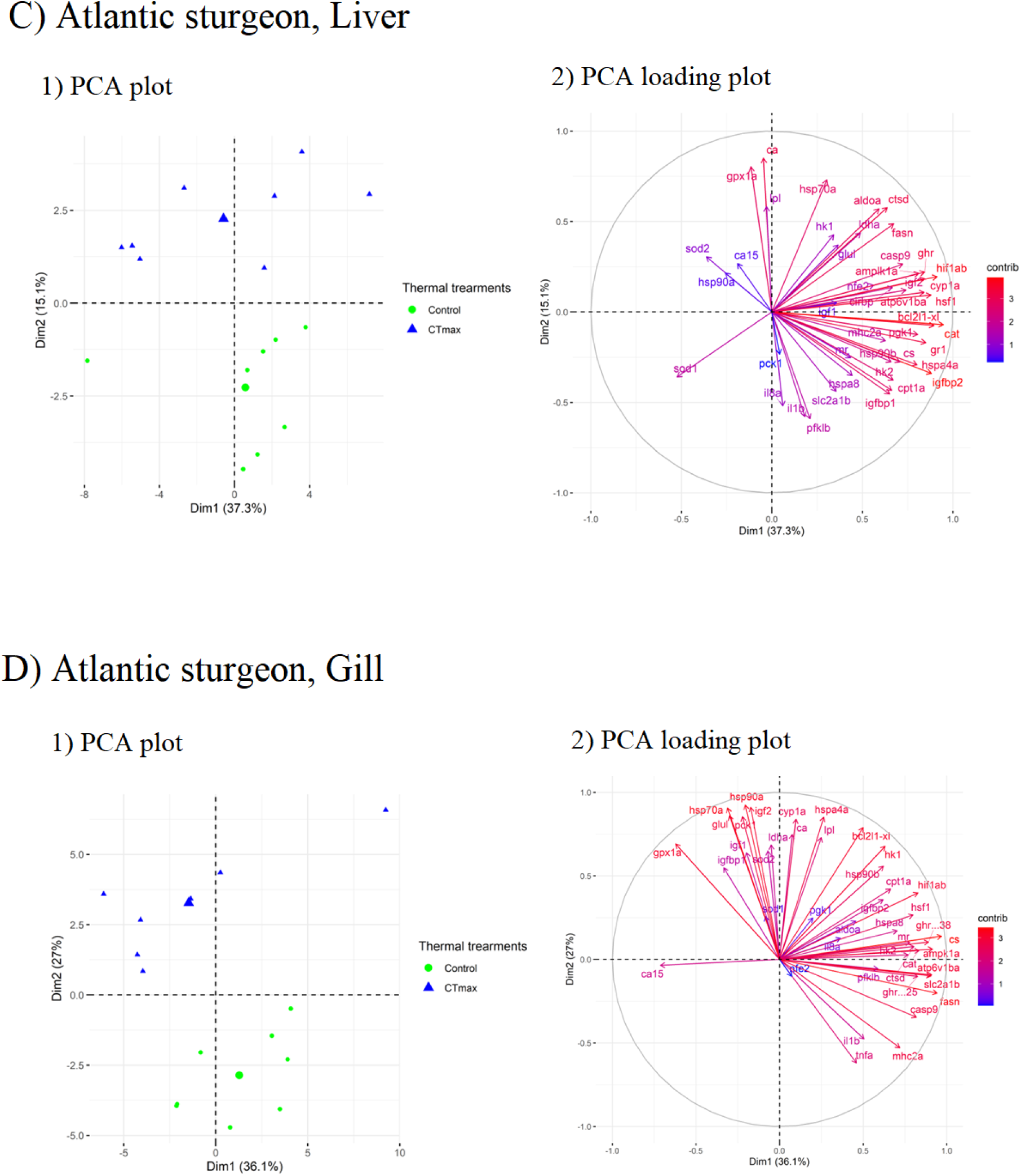
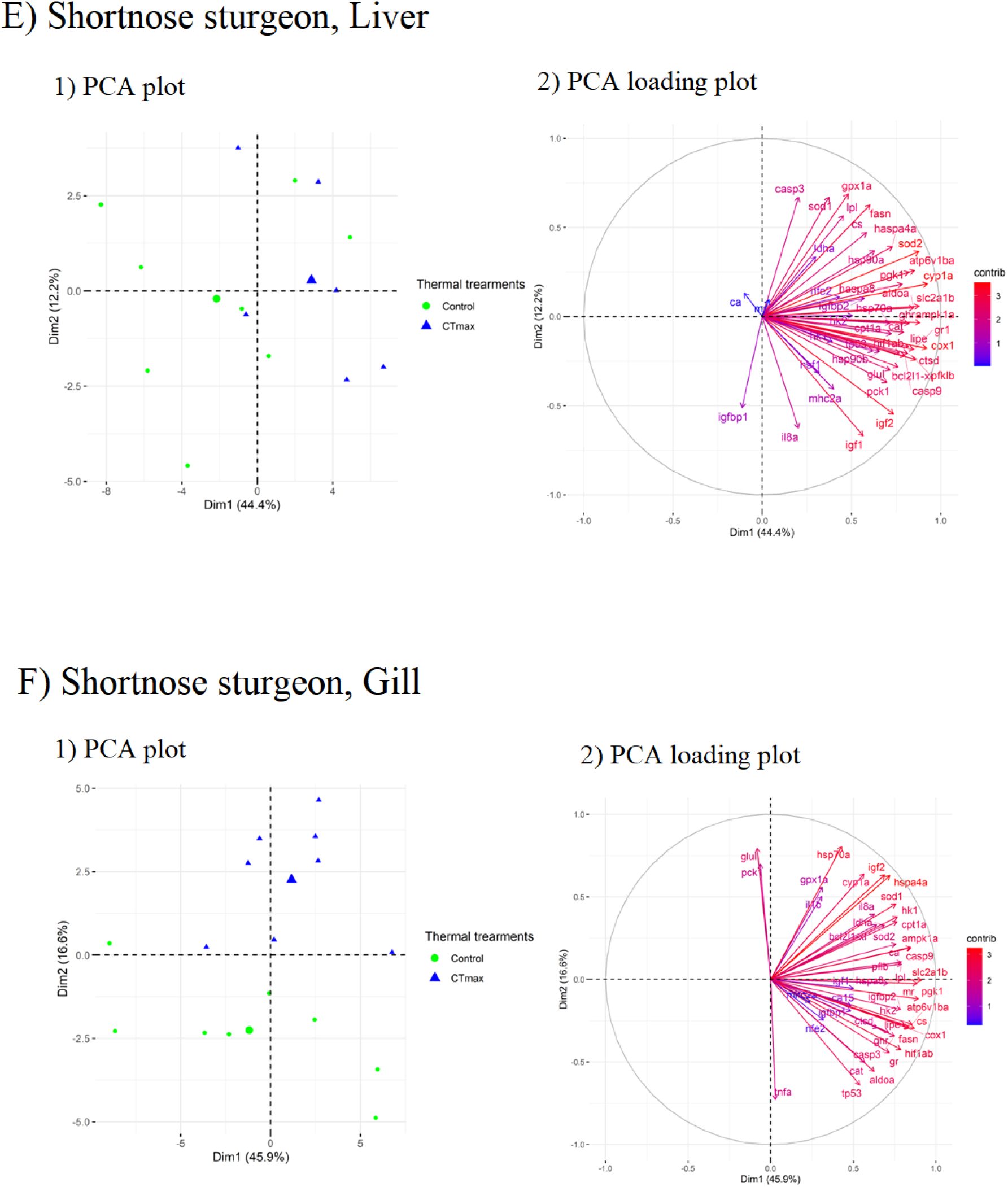
Plots from Principal Component Analysis showing the overall similarity in the expression pattern of 52 genes studied in three *Acipenser* species, and Loading score plots showing the contribution and direction of each gene in PC1 (Dim1) and PC2 (Dim2) in each species. Lake sturgeon were acclimated to 16, 20, and 24 °C and then experienced CT_max._ Atlantic and lake sturgeon were acclimated to 10 °C before CT_max_. Expression of each gene was measured in the samples collected before and after CT_max_ using qPCR. To generate the PCA plots, the C_t_ value of each gene was normalized to the mean C_t_ value of four control genes to generate unsupervised PCA plots.

## Discussion

The multispecies *Acipenser* OpenArray™ chip developed and presented in this study was able to amplify a wide selection of genes across multiple functional categories in three sturgeon species. Relative gene expression analysis showed that acute thermal stress changed the expression pattern of genes in the immune system, metabolism, osmoregulation, response to hypoxia, apoptosis, general stress response, and oxidative stress response. This expression pattern was affected by acclimation temperature, species, and tissues under study. Acute thermal stress affected the mRNA expression of 17-26 of the thermal metabolic stress genes that were assayed using the chip, depending on the species. Based on efficiency analysis, 52 genes in the lake sturgeon, 48 genes in the gill and liver of Atlantic sturgeon, 53 genes in the gill, and 52 genes in the liver of shortnose sturgeon were suitable for relative expression analysis. Across tissues and species, 46 of these genes were consistently suitable for analysis and can be readily applied to assess stressors for these sturgeon species. Future versions of this *Acipenser* chip will then need to re-design the assays with poor efficiency until there is a full suite of 56 assays that work across each species and tissue.

### Heat shock response and aerobic metabolism genes

Heat shock proteins are well-described markers of cellular stress, and their production is induced by a variety of factors including acute thermal stress (Sharp et al., 1999). Among the HSPs studied, *hsp70a* and *hsp90a* were significantly upregulated in response to acute thermal stress in all the species and tissues, except in the liver of the Atlantic sturgeon where *hsp90a* was downregulated. Upregulation of *hsp70a* and *hsp90a* is a common response to temperature and has been reported in several species (Shin et al., 2018; Sun et al., 2021; Yan et al., 2017). These two protein chaperones have been known to cooperate with the assistance of another protein called Hsp70/Hsp90 organizing protein, HOP, to regulate the folding of proteins such as hormone receptors (Odunuga et al., 2004). The upregulation of these two proteins has also been reported in response to increased temperature in sturgeons (Luo et al., 2022; Chen et al., 2023; Simide et al., 2016; Bugg et al., 2020; Penny et al., 2023), however, this upregulation was not significant in the muscle of shortnose sturgeon (Penny et al., 2023). Both hsp70 and hsp90 are inducible heat shock proteins, and their rapid upregulation has been reported in response to protein denaturation under thermal stress (Shin et al., 2018; Sun et al., 2021; Yan et al., 2017; Chen et al., 2005; Nguyen et al., 2021). The consistency in upregulation of *hsp70* and *hsp90a* in response to acute thermal stress across species and tissues makes them useful biomarkers for the detection of thermal stress in *Acipenser* sturgeons.

Genes such as *hk1, pgk1*, and *slc2a1b,* which are involved in the transportation of glucose to the cell and glycolysis (Augustin, 2010; Chandel, 2021), were downregulated in the gill tissue of the lake sturgeon in response to acute thermal stress. Downregulation of these genes may be a sign of decreased glycolysis. In addition, *pck1*, which plays a critical regulatory role in gluconeogenesis (Yu et al., 2021), was upregulated in both populations of lake sturgeons. This upregulation in *pck1* is also detected in the gill tissue of Atlantic and shortnose sturgeon, although genes involved in glycolysis were not downregulated. In the gill tissue of Atlantic sturgeon, the expression of *ldha* was also upregulated. The upregulation of *ldha* has been reported in other fish species in response to thermal stress (Cheng et al., 2018). Upregulation of *ldha* along with *pck1* may suggest that anaerobic glycolysis and gluconeogenesis are mechanisms induced simultaneously in response to acute thermal stress. However, in the gill tissue of shortnose sturgeon, *cpt1a* was upregulated. The protein coded by *cpt1a* is a key rate-limiting enzyme in fatty-acid oxidation. (Schlaepfer & Joshi, 2020). Increased expression of *cpt1a* can be an indicator of the higher mitochondrial oxidation of long-chain fatty acids. Changes in the expression of genes involved in the transportation of glucose, glycolysis, gluconeogenesis, and lipid oxidation after CT_max_ may be indicative of an interruption in the metabolism of energy caused by acute thermal stress.

Expression of *ampk1a* was upregulated in the liver of shortnose sturgeon. Because *ampk1a* works as an energy sensor in the cell and is activated by an increase in the ratio of AMP/ATP and/or ADP/ATP (Ke et al., 2018), its upregulation might be a sign of decreased production, or increased demand, for ATP during acute thermal stress. This upregulation was paralleled by an upregulation of *slc2a1b* and *ctsd,* which might be a cellular strategy to increase the import of glucose and the breakdown of proteins for energy production. Glucose is often stored as glycogen in the liver and can be released (Adeva-Andany et al., 2016) by glycogenolysis following hormonal stimulation under stressful conditions (Rabasa & Dickson, 2016). A decrease in the glycogen level in the liver, along with an increase in ATPase level, has been reported in aquatic species exposed to thermal stress (Han et al., 2023). In this study, these responses to acute thermal stress suggest an increase in the demand for ATP in shortnose sturgeon’s liver, which upregulates *ampk1a* to provide cells with energy-storing molecules by breaking down proteins, transporting glucose, and glycogenolysis.

### Oxidative stress response and apoptosis

Changes in the expression of genes with antioxidant properties such as *gpx*, *sod*, and *cat*, have been observed in several fishes following thermal stress (Roychowdhury et al., 2021; Liu et al., 2024). In the present study, the expression of *cyp1a* was upregulated in the gill tissue of all three species after CT_max_. This gene is important in the metabolism of polyaromatic hydrocarbons and is usually upregulated by activating the Aryl hydrocarbon receptors (Ah-R) (Collier & Pritsos, 2003), however, it is also involved in the production of anti-inflammatory molecules such as epoxyeicosatrienoic acid and helping to maintain normal physiological function through cell signaling and immunity (Stading et al., 2020). Upregulation of pro-inflammatory cytokines, such as interleukins, was observed in this research and also in similar studies (Liu et al., 2022; Zhao et al., 2024; Pérez-Casanova et al., 2008; Guo et al., 2023), which might be triggered by upregulation of *cyp1a*.

Although *glul* was upregulated in the gill tissue of Atlantic sturgeon, it showed significant downregulation in the gill tissue of Winnipeg River lake sturgeon acclimated to 24 °C. Glutamate-ammonia ligase, the protein coded by *glul*, is a glutamine synthetase important in detoxifying ammonia (Marginean et al., 2013; Ip & Chew, 2010). The upregulation of glutamine synthetases has been reported in response to increased ammonia concentrations in aquatic animals (Liu et al., 2014; Banerjee et al., 2018). Because increased temperature can increase the production and excretion of ammonia in fish (Zheng et al., 2008; Kieffer & Wakefield, 2009), upregulation of this gene might be a detoxification response to increased production and release of ammonia in Atlantic sturgeon. Downregulation of other oxidative stress response genes, such as *cat*, *gpx1a*, *nfe2*, *sod1*, and *sod2,* in the gill tissue of the Winnipeg River lake sturgeon may be an indicator of decreased detoxification potential of those cells under acute thermal stress.

However, the expression of *gpx1a* and *sod1* was upregulated in the gill tissue of the Atlantic sturgeon and shortnose sturgeon, respectively. This increase might be a response to an increase in the production of hydrogen peroxide or superoxide radicals, which have been reported in a variety of fish species (Chen et al., 2021; Liu et al., 2022; Yu et al., 2017; Dash et al., 2023). The expression of oxidative stress response genes did not show significant changes in the liver, showing the lower sensitivity of the oxidative stress response of this organ to acute thermal stress or lower production of ROS.

In lake sturgeon, acute thermal stress significantly downregulated the expression of pro-apoptotic genes *casp9* and *tp53,* although acclimation to higher temperatures resulted in a moderate upregulation of these genes after 30 days. Significant downregulation of *casp9* and upregulation of *bcl2l1-xl* was also observed in the gill tissue of Atlantic sturgeon. The protein coded by *casp9* has an important role in activating *tp53-*dependent apoptosis (Wu and Ding, 2002). However, *bcl2l1-xl* is an antiapoptotic protein that aids in cellular survival by regulating reactive oxygen species production (Loo et al., 2020). The downregulation of *casp9* and *tp53*, and the upregulation of *bcl2l1-xl* may be signs of decreased apoptosis during acute thermal stress.

### Immune response genes

Acclimation to higher temperatures has been reported to reduce the immune system response to pathogens in lake sturgeon (Bugg et al., 2023). However, acclimation to higher temperatures increased the expression of *mhc2a* in both populations of lake sturgeon, although this increase only happened at 20 °C acclimation temperature for the Burntwood River population. MHC class II is usually produced in the thymic epithelial cells or antigen-presenting cells, and other cells do not produce this protein unless they are stimulated by *ifng* caused by infection, inflammation, or trauma (Reith et al., 2005). Similarly, *il8* also showed an upregulation pattern similar to *mhc2a* in both populations. The upregulation of immune response genes such as *myd88*, *c3* and *lysozyme c* caused by acclimation to the higher temperature (20 °C) has been reported in lake sturgeon, however, fish acclimated to higher temperatures also had lower expression of *il1b* following exposure to bacterial lipopolysaccharides (LPS) (Bugg et al., 2023). Additionally, *il1b* showed significant downregulation in the Burntwood River population in response to acclimation to 24 °C. Interleukins are proteins with a pro-inflammatory role in animals (Secombes & Bird, 2011). The significant change in the expression of these genes, along with the upregulation of *mhc2a* in the gill, might be a sign of an inflammatory response in gill tissue to acute thermal stress.

Lake sturgeon exposed to acute thermal stress had a downregulation of *il1b* in the Burntwood River population and *il8* in the Winnipeg River population. Atlantic sturgeon showed significant downregulation of *il1b, tnfa,* and *mhc2a* in the gill, and *il1b* and *il8* in the liver. In shortnose sturgeon, although *tnfa* was significantly downregulated, *il1b* and *il8* were upregulated in the gill, and liver tissue did not show any significant change in the expression of genes involved in the immune system. Genes such as *il1b*, *il8*, and *tnfa* code for cytokines with important roles in the immune system, especially inflammation (Demir, 2022). These proteins are usually produced by white blood cells, or other non-immune cells, during inflammation and render their role by mechanisms such as activation of other white blood cells, induction of other cytokines, inhibition of neutrophil adhesion to the vascular endothelium, and stimulation of angiogenesis (Koper-Lenkiewicz et al., 2022). Differential expression patterns of these genes in sturgeons in this study can be because of different sensitivities of species to inflammation caused by thermal stress.

Together, these molecular responses demonstrate some differentiation in the response to thermal stress across sturgeon species, but also conserved mechanisms employed by diverse sturgeon species and populations to respond to this key stressor. The assays used to measure these responses are widely applicable across the species assessed here and can be used to evaluate the responses to thermal stress across different biological mechanisms. Gill-based assays can now be readily applied to non-lethally assess the response of sturgeons to acute thermal stress and provide new insights into the effects of this key stressor in future studies or the monitoring of threatened wild populations.

## Author contributions

KMJ and NJB were involved in funding acquisition, contributed to gene selection and developing the *Acipenser* OpenArray™ chip, and supervised the research. WSB designed and performed experiments on lake sturgeon. FMP and SAP designed and performed experiments on Atlantic and shortnose sturgeon. HH and KM designed and tested the primers and probes for the OpenArray™ chip. HH ran the OpenArray assays, analyzed the data, and wrote the first draft. All the authors read and provided comments on the manuscript.

## Acknowledgments

The authors thank the Genome Canada funded GEN-FISH project and an NSERC Discovery Grant to KMJ for funding this research. We thank Mark Shrimpton for providing white sturgeon samples for testing primers. We also thank the staff of the Animal Holding Facility at the University of Manitoba and the University of New Brunswick.

